# Human brown adipose tissue demonstrates substantial ^18^F-fluorocholine uptake at room temperature for phosphatidylcholine synthesis

**DOI:** 10.1101/2025.08.24.671970

**Authors:** Karla J Suchacki, Lynne E Ramage, Luke D Boyle, T’ng Choong Kwok, Giovanny Rodriguez Blanco, Alexandra Kelman, Calum D Gray, Alex von Kriegsheim, Maria-lena Gregoriades, Gabriel C Oniscu, Alison M Fletcher, Natalie Z M Homer, John D Terrace, J. William Allwood, Sonia J Wakelin, Edwin JR van Beek, Dilip Patel, Andrew J Finch, Roland H Stimson

## Abstract

**Objective:** Brown Adipose Tissue (BAT) is composed of mitochondrial-rich, multilocular adipocytes, which dissipate energy to produce heat. Quantification of BAT mass is most commonly performed by ^18^F-fluorodeoxyglucose positron emission tomography (PET), which requires antecedent cold exposure. We hypothesized that ^18^F-fluorocholine PET could detect human BAT due to its requirement for considerable phosphatidylcholine synthesis, secondary to brown adipocytes’ high mitochondrial density and multiple lipid droplets.

**Methods:** 1) Six healthy men with detectable ^18^F-fluorodeoxyglucose uptake by BAT were recruited to a randomised crossover study investigating ^18^F-fluorocholine uptake by BAT during warm and cold exposure. 2) ^18^F-fluorocholine uptake by supraclavicular adipose tissue was quantified in 76 patients who had undergone ^18^F-fluorocholine PET/CT scanning. 3) Choline transporter expression was quantified in human BAT and white adipose tissue (WAT), and in brown and white adipocytes. 4) Lipidomics was performed on human brown adipocytes incubated with ^15^N-choline to determine phospholipid synthesis.

**Results:** 1) ^18^F-Fluorocholine uptake by BAT was substantially greater than by WAT during both warm and cold exposure. 2) ^18^F-Fluorocholine uptake by supraclavicular adipose tissue was higher in the colder seasons, inversely associated with body mass index, and positively associated with tissue radiodensity. 3) Expression of the choline transporter *SLC44A3* was higher in human supraclavicular BAT than WAT, while *SLC44A2* was expressed more highly in human brown than white adipocytes. 4) ^15^N-choline tracing in human brown adipocytes identified incorporation into >30 phosphatidylcholine species.

**Conclusions:** ^18^F-fluorocholine PET may be a novel method to quantify human BAT at room temperature.

**Highlights:**

- ^18^F-Fluorocholine (FCH) PET can detect human BAT without the need for cold exposure.
- ^18^F-FCH uptake by supraclavicular AT is inversely associated with body mass index.
- The choline transporter SLC44A3 is more highly expressed in BAT than WAT.
- Human brown adipocytes utilise choline for phosphatidylcholine synthesis.

GRAPHICAL ABSTRACT

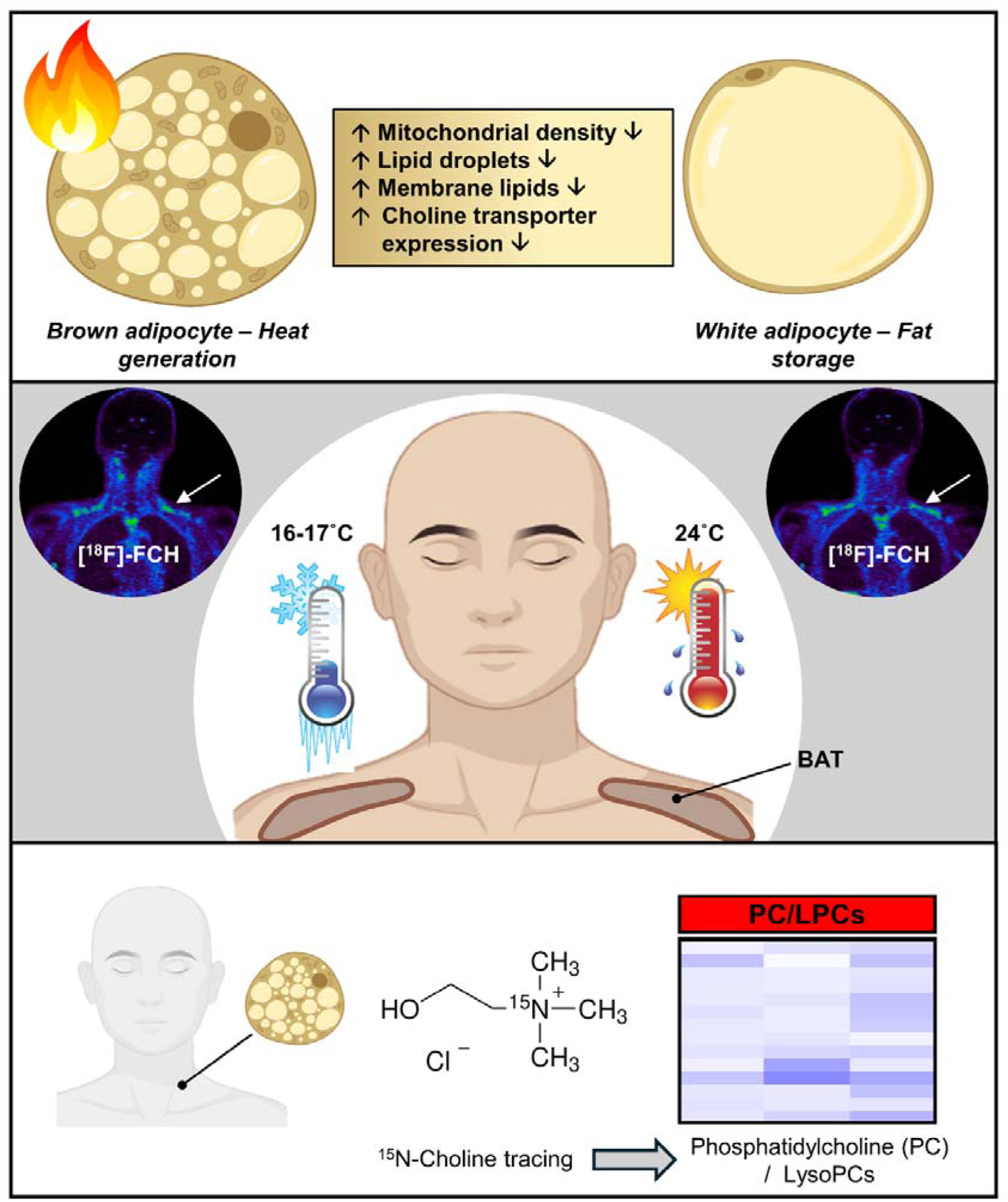

## 1. INTRODUCTION

Energy balance is the relationship between energy input and energy output. Obesity occurs when energy intake chronically exceeds energy expenditure, over the past 30 years the global prevalence of obesity has increased dramatically with now over a billion people in the world living with obesity, approximately 1 in 8 of the global population [1]. Obesity is associated with multiple co-morbidities such as type 2 diabetes mellitus (T2DM), dyslipidaemia, hypertension, certain cancers, and cardiovascular disease (CVD), resulting in increased morbidity and mortality [2]. Weight reduction through lifestyle modifications such as limiting food intake and increasing physical activity is difficult to maintain, hence there is considerable interest in pharmacological approaches. The recent development of agents such as glucagon-like peptide-1 agonists and dual agonists can affect 10-20% weight loss through reduction in appetite and food intake [3, 4], but weight regain occurs following treatment withdrawal, weight loss is considerably less in those with T2DM [5], and many health boards do not have the resources to offer these agents to most patients with obesity at present, as such the prevalence of obesity will likely continue to rise in the short term. In addition, weight loss induces a compensatory reduction in energy expenditure, which limits further weight reduction [6]. Therefore, pharmacotherapy to increase energy expenditure is another attractive therapeutic target, potentially in combination with appetite suppressants.

The relatively recent discovery of brown adipose tissue (BAT) in adult humans has garnered greater interest in this therapeutic approach, as the primary role of BAT is to generate heat to maintain body temperature in a cold environment [7]. Unlike white adipose tissue (WAT), brown adipocytes contain multiple small lipid droplets and many mitochondria which are equipped with a specialised thermogenic protein called uncoupling protein 1 (UCP1) [8]. Upon cold stimulus, sympathetic neurons innervating BAT release noradrenaline (NADR) from the synapse which activates various β-adrenergic receptors on the brown adipocyte, stimulates lipolysis and activates UCP1, uncoupling respiration from ATP synthesis. BAT thermogenesis also induces substantial glucose uptake by BAT, which is exploited by the use of ^18^F-fluorodeoxyglucose positron emission tomography (^18^F-FDG-PET) to quantify BAT mass and activity [9, 10]. Highlighting its potential importance, individuals with detectable ^18^F-FDG uptake by BAT or with high UCP1 expression in supraclavicular AT biopsies have a lower prevalence of cardiometabolic risk factors and cardiovascular disease [11, 12]. However, only ∼5% of patients have detectable ^18^F-FDG uptake by BAT at room temperature, so active cooling of participants is necessary in order to determine whether people have BAT [9, 13, 14]. In addition, there are no circulating biomarkers to reliably identify individuals with and without BAT. Therefore, PET radiotracers capable of detecting BAT without cold exposure would be of substantial benefit, which can also provide important physiological insights into BAT function. Specialised radiotracers such as those binding to the norepinephrine transporter can detect BAT [15], but these are not widely available. However, there are no known PET radiotracers in routine clinical use that can identify BAT without cooling.

Phospholipids such as phosphatidylcholines (PCs) are key components of the membranes of lipid droplets and mitochondria, in addition to surface cell membranes [16]. PCs also play an important role in maintaining smaller lipid droplet size [17], a hallmark of multiloculated brown adipocytes. Choline containing phospholipids such as PCs and lysoPCs (LPCs) are highly abundant in BAT compared with WAT depots in mice, and cold exposure increases the abundance of specific PCs in BAT [18, 19]. PCs are synthesized from choline [20], and we hypothesized that enhanced requirements for PC synthesis would result in greater choline uptake by BAT than WAT, even during warm conditions. ^18^F-Fluorocholine (^18^F-FCH) is a PET radiotracer commonly used in the investigation of prostate cancer and hyperparathyroidism so is widely available [21, 22]. Here we report that human supraclavicular BAT demonstrates substantial [^18^F]-FCH uptake during warm conditions, which increases further during cold exposure. In addition, ^18^F-FCH uptake by BAT was inversely associated with BMI in a retrospective analysis of clinical scans performed at room temperature. Finally, we have demonstrated greater expression of the choline transporter *SLC44A3* in human BAT than WAT, greater *SLC44A2* expression in brown adipocytes than white adipocytes, and identify the PC species formed in human brown adipocytes using labelled choline.

## 2. RESULTS

### 2.1 Human BAT demonstrates substantially greater ^18^F-FCH uptake during warm conditions than WAT

To determine whether human BAT can be detected by ^18^F-FCH PET, six healthy young adult male volunteers (mean ± SD, aged 22.0 ± 3.3 years) of normal weight (body mass index (BMI) 22.8 ± 1.9 kg/m^2^) were recruited to a randomised crossover study (Figure 1A). Additional baseline characteristics have been reported previously [12]. At visit 1, participants were exposed to 2h of mild cold exposure at (mean room temperature 16.1 ± 0.8°C) to activate BAT, then underwent an ^18^F-FDG PET/MRI scan to quantify their BAT. The ^18^F-FDG and anthropometric data from these participants have been presented previously [12]. ^18^F-FDG uptake by BAT was detected in all volunteers, confirming these subjects had active BAT during acute cold exposure (Figure 1B-D). Following confirmation of detectable BAT, the volunteers proceeded to the final 2 study visits.

**Figure 1.**
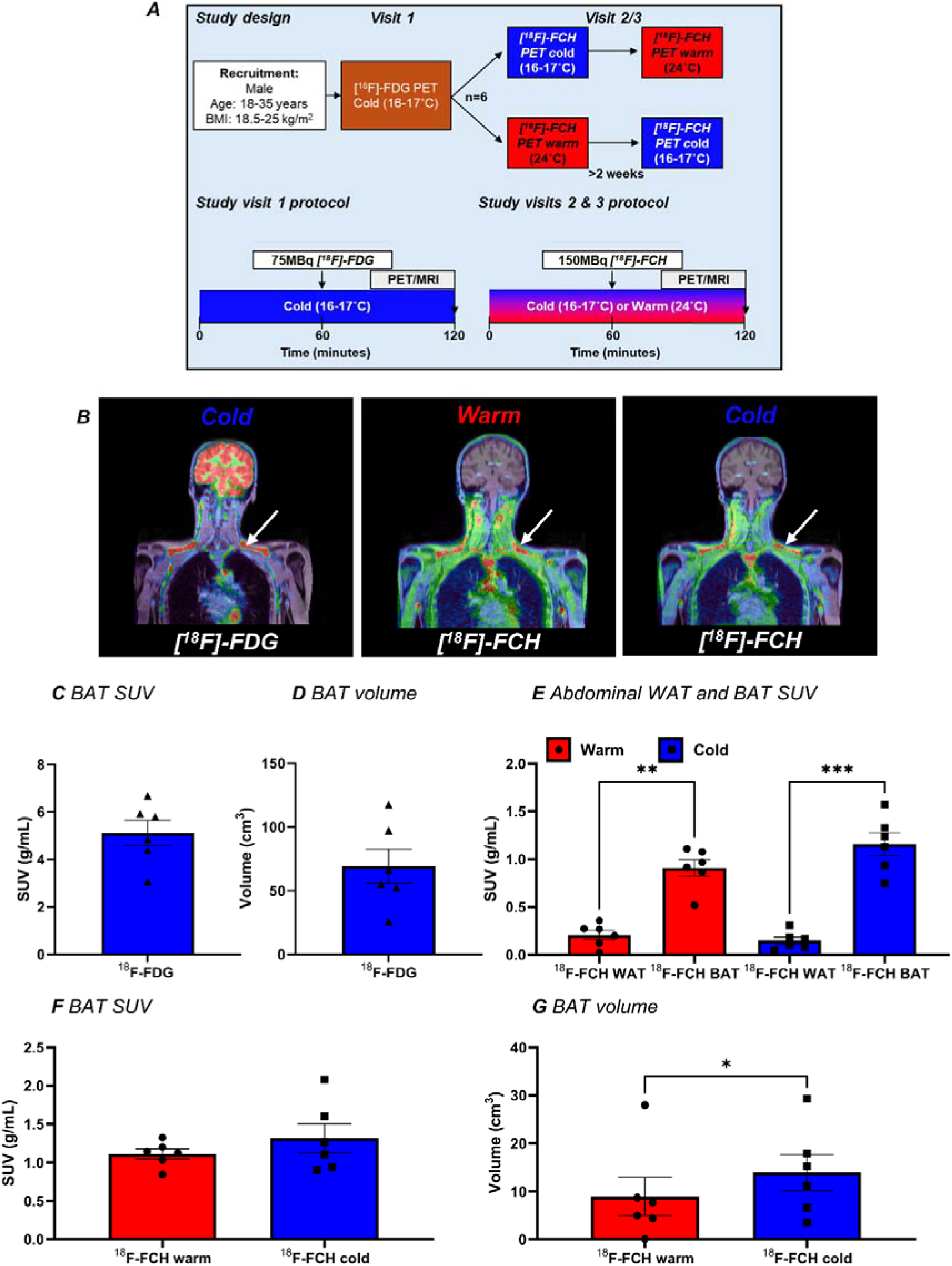
^18^F-FCH uptake by BAT in healthy volunteers during warm and cold exposure. A) Six normal weight healthy men were recruited to a randomised crossover study. Volunteers initially underwent an ^18^F-FDG PET/MRI scan following mild cold exposure, then attended on 2 further occasions for ^18^F-FCH PET/MR scans during either warm or cold conditions. B) Representative fused PET/MR images following 75MBq ^18^F-FDG (left panel) and ^18^F-FCH during warm (middle panel) and cold exposure (right panel). C) ^18^F-FDG uptake by BAT and D) BAT mass quantified by ^18^F-FDG in all volunteers. E) ^18^F-FCH uptake by BAT quantified using the ^18^F-FDG ROIs, and abdominal WAT during warm (red columns) and cold (blue columns) exposure. F) ^18^F-FCH uptake by BAT and G) BAT mass quantified using a standardised uptake value threshold of ≥0.5 g/mL. Data are represented as mean ± SEM, and were analysed using repeated measures one-way ANOVA with Tukey’s multiple comparisons tests (E), paired t-test (F), Wilcoxon test (G). Significant P values are detailed in the respective panels. *p<0.05, **p<0.01, and ***p<0.001.

At visits 2&3, volunteers underwent ^18^F-FCH-PET/MR scanning following 2h of either warm (room temperature 24.3 ± 1.2°C) or cold (16.2 ± 0.5°C, P<0.001 vs warm conditions) exposure in random order. Initially, regions of interest (ROIs) identified from the ^18^F-FDG scans were co-registered to the ^18^F-FCH images to quantify ^18^F-FCH uptake by BAT. In contrast to ^18^F-FDG which does not demonstrate substantial uptake by BAT during warm conditions in most participants [14], there was detectable ^18^F-FCH uptake by BAT during warm exposure in all volunteers, and ^18^F-FCH uptake was similar between warm and cold visits (Figure 1B/E). Importantly, ^18^F-FCH uptake by BAT was ∼4-fold higher than in abdominal WAT (Figure 1E). Mean ^18^F-FCH uptake was >0.5 g/mL in BAT and was <0.5 g/mL in WAT all volunteers, suggesting an ^18^F-FCH threshold could be employed to identify BAT without prior ^18^F-FDG-PET during cold exposure. Therefore, we generated ROIs in these ^18^F-FCH-PET/MR scans using a SUV threshold of ≥0.5 g/mL, in addition to a fat fraction threshold of 50% to remove non-adipose tissue depots as described previously [23]. Using this technique, ^18^F-FCH uptake successfully detected supraclavicular BAT in all subjects during warm exposure, demonstrating a possible role for ^18^F-FCH PET in the identification of supraclavicular BAT (Figure 1F). In addition, cold exposure increased the volume of detectable BAT using ^18^F-FCH by ∼50%, but BAT SUV and volume were lower using ^18^F-FCH than using ^18^F-FDG during cold exposure (Figure 1F/G). However, the 0.5 g/mL threshold did not detect other BAT depots typically identified by ^18^F-FDG such as paraspinal BAT, highlighting possible depot-specific differences. In addition, no other WAT depots were identified using the 0.5 g/ml threshold, confirming higher uptake by BAT. Compared with warm conditions, cold exposure increased circulating noradrenaline and non-esterified fatty acid concentrations consistent with activation of the sympathetic nervous system and lipolysis [24, 25], without altering glucose or insulin (Figure S1).

### 2.2 ^18^F-FCH uptake by supraclavicular adipose tissue at room temperature is inversely associated with BMI in patients

Given that ^18^F-FCH could detect supraclavicular BAT in normal weight young adult volunteers without prior cold exposure, we undertook a retrospective analysis of 76 patients with prostate cancer or primary hyperparathyroidism who had undergone ^18^F-FCH-PET/CT scanning at room temperature as part of their standard clinical care (Figure 2A, Table S1). Previous studies using ^18^F-FDG-PET and measuring UCP1 expression in BAT have demonstrated that these are inversely associated with age and BMI, in keeping with reduced BAT activity in older obese individuals [9, 11, 12]. In addition, BAT is associated with greater radiodensity on CT scanning (as measured using Hounsfield units or HU) than WAT [26]. Therefore, we quantified ^18^F-FCH uptake in supraclavicular adipose tissue (AT) (the most typical location for BAT) in these patients to determine any associations with these factors. The patients included in this retrospective analysis were significantly older (mean age 59 ± 14 years) and heavier (BMI 28 ± 5 Kg/m^2^) than those who participated in the healthy volunteer study. ^18^F-FCH uptake by supraclavicular AT was associated positively with tissue radiodensity, and inversely associated with BMI (Figure 2B-C). However, ^18^F-FCH uptake was not associated with age in this cohort (Figure 2D). ^18^F-FDG data has demonstrated that BAT activity is greater in the colder months [9, 11], so we also tested whether ^18^F-FCH uptake showed seasonal change. ^18^F-FCH uptake by supraclavicular adipose tissue was greater in the winter/spring than in the summer/ autumn seasons (Figure 2E). These data demonstrate that ^18^F-FCH uptake by supraclavicular AT at room temperature is a marker of human BAT.

**Figure 2.**
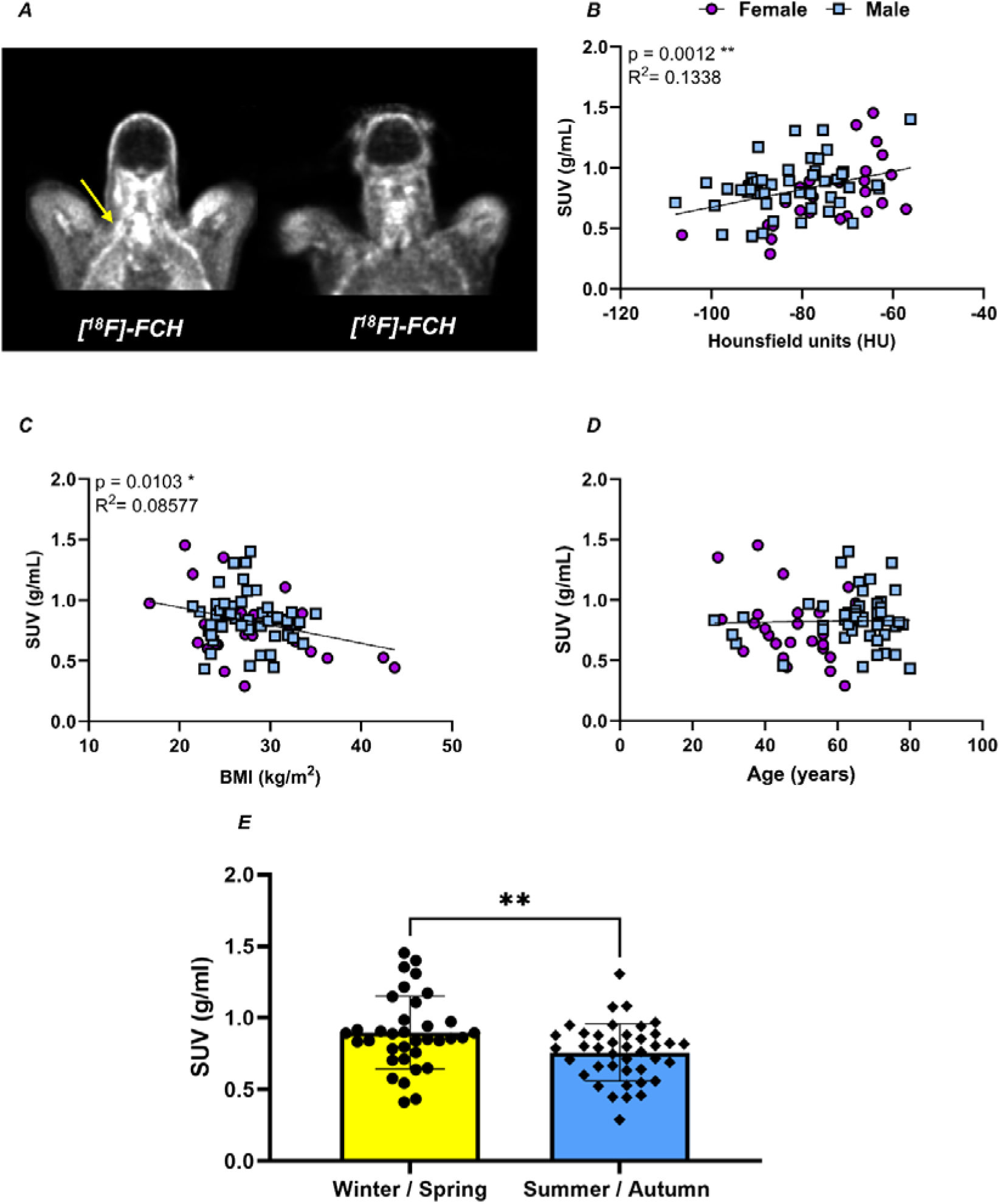
^18^F-FCH uptake by supraclavicular adipose tissue in patient scans. A) Representative fused PET/CT scans from a patient with high (left image) and low (right image) ^18^F-FCH uptake by supraclavicular adipose tissue. B-D) ^18^F-FCH uptake by supraclavicular adipose tissue was quantified in 26 female (purple circles) and 50 male (blue squares) who had undergone PET/CT scanning as part of their standard care for either prostate cancer (n=41) or primary hyperparathyroidism (n=35). ^18^F-FCH uptake was B) positively associated with supraclavicular adipose tissue radiodensity, and C) negatively associated with body mass index, but D) was not associated with age. E) ^18^F-FCH uptake was higher in the colder (yellow columns, Dec-May) vs warmer seasons (blue columns, Jun-Nov). Data were analysed by A-D) simple linear regression or E) unpaired t tests. Significant P values are detailed in the respective panels. *p<0.05, **p<0.01.

### 2.3 *SLC44A3* expression is higher in human supraclavicular BAT than WAT

Choline enters cells primarily through the high affinity transporter *SLC5A7*, intermediate affinity choline-like transporters *SLC44A1-5*, and the low affinity organic cation transporters *SLC22A1/2 [16]*. In addition, the inner mitochondrial membrane choline transporter *SLC25A48* plays a role in BAT thermogenesis [27]. To determine whether expression of these transporters was increased in BAT, we analysed mRNA expression levels in previously published transcriptomic data from primary human brown and white adipocytes [23]. There was substantial expression of *SLC44A1-3* in both brown and white adipocytes but not the other transporters (Figure 3A), qPCR confirmed higher *SLC44A2* expression in human supraclavicular brown than white adipocytes (Figure 3B). In whole tissue, the *SLC44A1-*3 transporters were highly expressed in BAT and WAT, and *SLC44A3* expression was higher in human supraclavicular BAT than WAT (Figure 3C), consistent with the enhanced ^18^F-FCH uptake observed *in vivo*. In both cells and whole tissue, *SLC22A1* demonstrated low expression, while *SLC22A2* was not detected. *SLC25A48* was only expressed in whole tissue, and expression was lower in BAT than WAT (Figure 3C). UCP1 expression was substantially greater in brown adipocytes and BAT than in white adipocytes and WAT respectively (Figure 3B/C). Choline transporter expression was also quantified in human peri-renal AT, a depot which did not demonstrate substantial ^18^F-FCH uptake *in vivo*. Previous data have revealed high UCP1 expression in AT obtained from near the upper pole of the kidney, the key marker of BAT [28]. Therefore, expression was quantified in AT obtained from the upper and lower poles of the kidney, along with subcutaneous abdominal WAT. Unlike in supraclavicular BAT, expression of *SLC44A3* was similar in peri-renal AT to abdominal WAT, despite high UCP1 expression in the upper pole (Figure 3D). These data suggest that enhanced choline uptake by supraclavicular BAT may be mediated via the SLC44A2/3 choline transporters.

**Figure 3.**
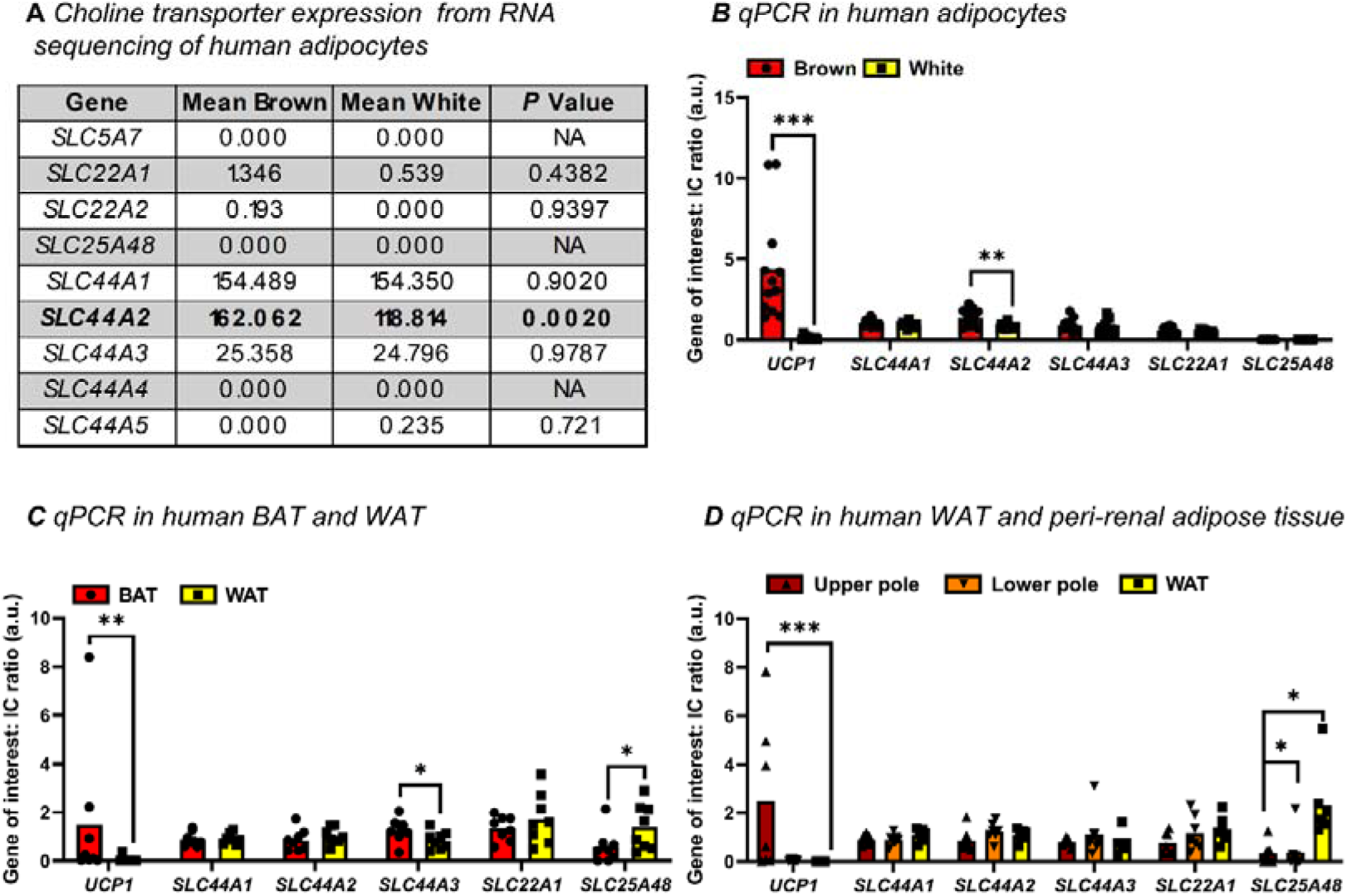
Choline transporter expression in human adipocytes and whole adipose tissue. A) Choline transporter expression from RNA sequencing of paired primary human brown and white adipocytes (n=4/ group), SLC44A2 is highlighted in bold as was more highly expressed in brown adipocytes. B-D) Data are mean ± SEM for qPCR of choline transporters in human adipocytes and whole adipose tissues. B) Choline transporter mRNA levels in paired primary human brown (red columns) and white adipocytes (yellow columns, n=12/ group), SLC44A2 expression was higher in brown adipocytes. C) mRNA levels in paired supraclavicular BAT (red columns) and neck WAT (yellow columns, n=8/group). SLC44A3 expression was higher in BAT, while SLC25A48 was lower in BAT. D) mRNA levels in paired peri-renal adipose tissue obtained from the upper pole (red columns), lower pole (orange columns), and subcutaneous abdominal WAT (yellow columns, n=7/ group). SLC25A48 expression was highest in WAT. Data were analysed using A) DESeq2 [23], B/C) paired t test or Wilcoxon matched-pairs signed rank and D) repeated measures ANOVA (Friedmann’s with Dunn’s or Holm-Sidak post-hoc tests).

### 2.4 Choline is converted into multiple PC species in human brown adipocytes

Following uptake of choline, PCs and LPCs are primarily synthesized through the Kennedy pathway and Lands cycle, [20, 29, 30], but may also be synthesized alternatively through the betaine pathway [31, 32] (Figure 4A). We first analysed existing RNA sequencing data from human brown and white adipocytes [23], and determined high transcript levels of the various enzymes involved in these pathways in both brown and white adipocytes, in keeping with utilisation of choline for PC synthesis (Figure 4A, Table S2). Of these genes, the phospholipase A2 *PLA2G4A* was expressed more highly in brown than white adipocytes, which is a key regulator of lipid droplet formation [33]. To determine the PC and LPC species synthesized from choline, we cultured human primary brown adipocytes either with or without [^15^N]-choline in the presence or absence of noradrenaline for 24 hours. Lipidomics was performed on the cell lysates to determine the lipids synthesized from (m+1)-choline. ^15^N-Choline was detected in >30 PC and LPC species in human brown adipocytes (Figures 4B and S2, Table S3), including PC 34:1, PC 34:2, PC 38:2, and LPC 16:0 which have previously been identified in murine BAT and/ or associated with cold-induced thermogenesis. However, noradrenaline treatment did not alter the abundance of ^15^N-containing lipid species in these cells. These data confirm that choline is utilised for PC/ LPC synthesis in human brown adipocytes.

**Figure 4.**
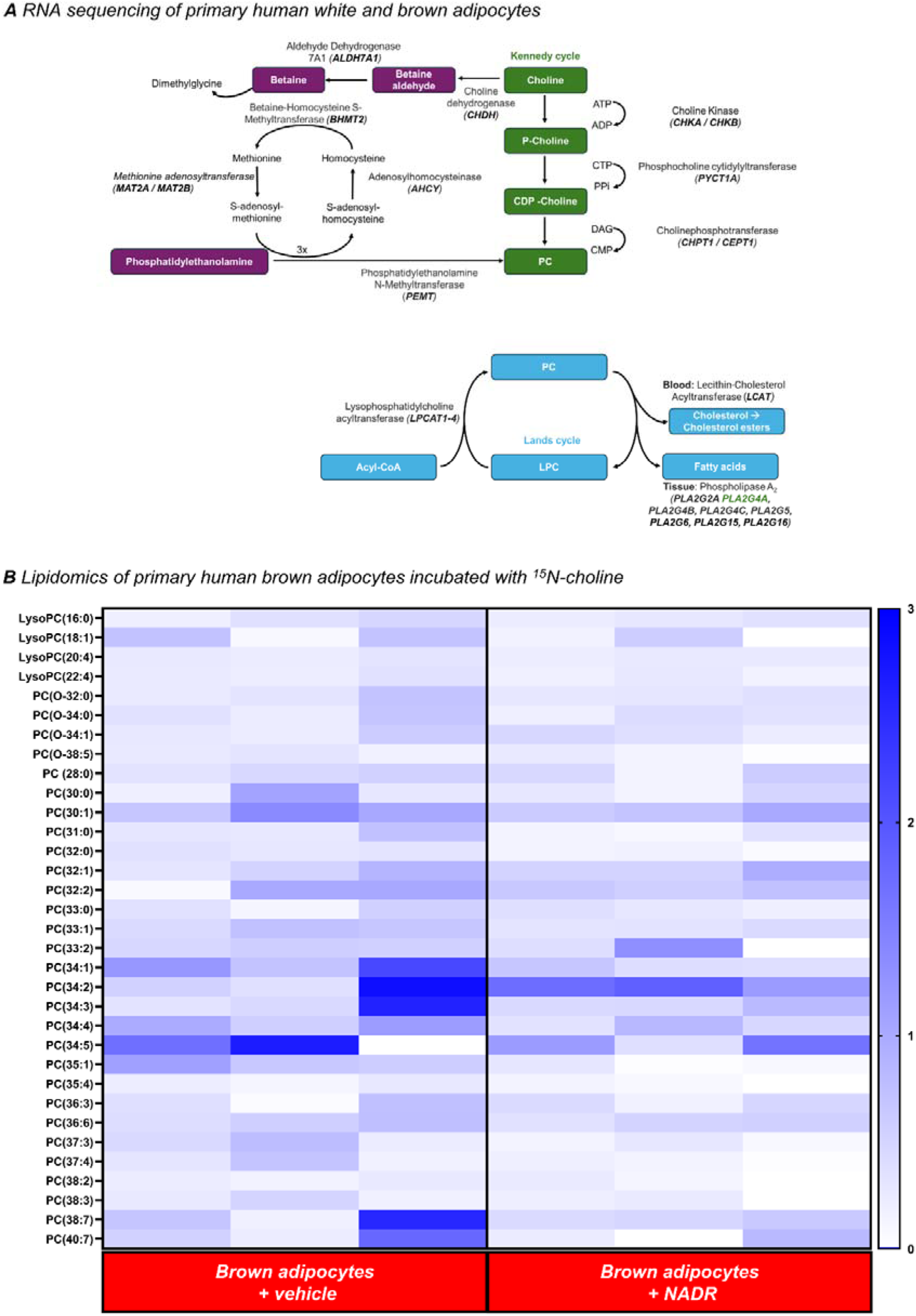
Choline is converted into multiple phosphatidylcholine species in human brown adipocytes. *A)* Choline can be metabolised through the Kennedy, Lands, and betaine pathways. Key genes in those pathways were expressed, as identified from RNA sequencing of paired primary human white and brown adipocytes (n=4/ group). Gene names written in black = expressed similarly in both cell types; green font = higher in brown adipocytes; no gene was more highly expressed in white adipocytes. B) Lipidomics was performed on human brown adipocytes incubated with either 400µM ^15^N-choline/ vehicle, in the presence or absence of 10µM noradrenaline (NADR) or vehicle for 24 hours. Heat map detailing the phosphatidylcholine (PC) and lyso-PC (LPC) species incorporating ^15^N-choline in vehicle and NADR-treated cells. Values presented are the ratios of (m+1)/(m+0), following theoretical subtraction of the contribution of (m+1) from the natural abundance of ^13^C. Data were analysed by Wilcoxon signed-rank test.

## 3. DISCUSSION

These data demonstrate, to our knowledge for the first time, that that the radiotracer ^18^F-FCH can be used to detect human BAT during warm exposure, in support of our hypothesis. In healthy volunteers, ^18^F-FCH uptake by supraclavicular AT during warm or cold exposure co-localised with ^18^F-FDG uptake by BAT during cold exposure. Rodent brown adipocytes also demonstrate a greater choline content than white adipocytes, as revealed using ^13^C-magentic resonance spectroscopy [34]. Importantly, ^18^F-FCH uptake was substantially greater in ^18^F-FDG-postive AT (BAT) than in AT without substantial ^18^F-FDG uptake (WAT) in these volunteers. Specific criteria have been proposed for SUV thresholds for static ^18^F-FDG PET imaging to detect human BAT [13], which we utilised in this current study to generate the BAT ROIs for comparison with abdominal WAT which demonstrated minimal ^18^F-FDG uptake. The mean ^18^F-FCH SUV for abdominal WAT in each of the six participants was <0.4 g/mL during warm and cold exposure, and was >0.5 g/mL in BAT, so an SUV threshold of 0.5 g/mL, in combination with a fat fraction consistent with adipose tissue, could reliably detect supraclavicular BAT in this young cohort. However, BAT mass identified by ^18^F-FCH using the 0.5 g/mL threshold was considerably lower than that quantified by ^18^F-FDG, suggesting that ^18^F-FCH PET is not as sensitive as ^18^F-FDG. Importantly, no other WAT depots were identified using the 0.5 g/mL ^18^F-FCH threshold, highlighting that ^18^F-FCH uptake by BAT was greater than all WAT depots.

We also investigated ^18^F-FCH uptake by supraclavicular AT in an older cohort with prostate cancer and primary hyperparathyroidism. In this group, ^18^F-FCH uptake was closely correlated with adipose tissue radiodensity, negatively correlated with BMI, and uptake was higher in the colder months in agreement with previous BAT data using ^18^F-FDG [9, 26]. These data suggest that ^18^F-FCH may be a widely available clinical PET tracer that can detect BAT without the need for prior cold exposure, and provide complementary data that BAT mass is decreased in obesity. However, in the older cohort the mean ^18^F-FCH SUV was >0.5 g/mL in the majority of patients, suggesting that this SUV threshold would not be a reliable discriminator of BAT in this cohort. Therefore, future studies combining ^18^F-FDG and ^18^F-FCH scanning will be required to assess generalisability in larger, more diverse cohorts, identify a suitable SUV threshold to discriminate BAT from WAT and determine the sensitivity and specificity of this technique.

The high choline uptake by BAT is most likely due to the increased requirement for membrane lipid synthesis due to the substantial mitochondrial content and multiple lipid droplets in this depot [35]. This conclusion is strengthened by the lack of a significant increase in the SUV of ^18^F-FCH uptake by BAT during cold exposure, which suggests that increased BAT perfusion does not drive the enhanced ^18^F-FCH uptake, as BAT perfusion doubles during cold exposure [25, 36]. However, cold exposure did increase the mass of detectable BAT using ^18^F-FCH when using the 0.5 g/mL threshold, which could be in keeping with increased utilisation of choline during human BAT thermogenesis. In support of these data, cold exposure increased synthesis of multiple PC (and phosphatidylethanolamine) species in murine BAT [37]. In addition, adrenergic stimulation of murine brown adipocytes enhanced lyso-PC synthesis which increased proton leak by UCP1, suggesting that increased PC synthesis may directly increase BAT thermogenesis [38]. However, noradrenaline did not alter the incorporation of ^15^N-choline into PC or LPC species in our human primary brown adipocytes over 24 hours. Recent rodent data have revealed that choline can enhance thermogenesis via pathways other than PC synthesis. For example, choline transport into BAT mitochondria is important for conversion to betaine which enhances BAT thermogenesis [27]. Although the mitochondrial choline transporter SLC25A48 was not expressed in our human brown adipocytes, the SLC44A transporter family are expressed in mitochondrial membranes [39, 40], so may be driving mitochondrial choline uptake in human BAT. In this current protocol, we only undertook lipidomic analyses of the primary adipocytes so were unable to determine whether ^15^N-choline was utilised for other pathways involving polar metabolites such as betaine synthesis.

We found increased expression of the choline transporter *SLC44A2* in human brown vs white adipocytes, and *SLC44A3* in supraclavicular BAT compared with WAT, which may drive the higher relative ^18^F-FCH uptake by BAT *in vivo*. Interestingly, we noted that ^18^F-FCH uptake by supraclavicular BAT was substantially greater than WAT, while other BAT depots such as para-spinal or peri-renal BAT which are commonly identified by ^18^F-FDG were not detectable by ^18^F-FCH [41]. In addition, choline transporter expression was not increased in peri-renal BAT compared with WAT, potentially in keeping with this finding. Importantly, these data reveal differences between BAT depots *in vivo*, in support of previous research in human biopsies suggesting that peri-renal BAT may be less active than the supraclavicular depot [28]. Molecular signatures can vary between BAT depots [42], and it is possible that distinct depots differ in their roles.

^15^N-Choline was incorporated into >30 PC/ LPC species in human brown adipocytes, several of which have been shown to be important to BAT function or altered by cold exposure in mice. ^15^N-containing phospholipid species synthesized by human brown adipocytes included LPC 16:0, previous data found that circulating concentrations of LPC 16:0 were positively associated with ^18^F-FDG uptake by BAT during cold exposure [43]. In addition, LPC 16:0 is increased in murine brown adipocytes following adrenergic stimulation, and enhanced UCP1-mediated uncoupled respiration [38]. Other unmeasured phospholipids such as phosphatidylethanolamines can also enhance UCP1-dependent respiration [44]. We found substantial incorporation of ^15^N-choline into PCs with 32-34 carbons such as 32:1, consistent with studies in murine brown and beige adipocytes [38]. Choline was also incorporated into PC 34:2, which is increased in rodent BAT following cold exposure [37]. Interestingly, acute cold exposure increased circulating PC 34:2 (and PC 34:1) concentrations in humans, consistent with enhanced PC synthesis [45]. We also found choline incorporation into PC 38:2, which is highly abundant in murine BAT mitochondria and increased by cold exposure [46].

There are specific limitations of this study, in addition to those described above. Firstly, we quantified ^18^F-FCH uptake in healthy volunteers all with detectable ^18^F-FDG uptake by BAT, while our NHS cohort patients had not undergone ^18^F-FDG scanning, so were unable to determine ^18^F-FCH uptake by supraclavicular AT in ‘BAT-negative’ individuals. Quantifying supraclavicular ^18^F-FCH uptake in individuals without substantial ^18^F-FDG uptake in this depot would be interesting, however ^11^C-acetate data have suggested that ^18^F-FDG PET may underestimate BAT activity in older cohorts [47]. Furthermore, our lipidomic analyses were not optimised for the quantification of non-choline containing phospholipids such as phosphatidylethanolamines, and future work will be required to determine if noradrenaline stimulation enhances synthesis of other species. Finally, we could not determine the role and importance of the individual PC species to human brown adipocyte function in this current work, and it will be important to determine if any enhance thermogenesis in future research.

In summary, we have adapted ^18^F-FCH-PET as a novel technique to detect supraclavicular BAT in humans without the need for cold exposure, exploiting the higher membrane synthesis requirements of BAT due to its higher mitochondrial content and multiple lipid droplets compared with WAT. We have also identified that ^18^F-FCH uptake differs between BAT depots, highlighting possible functional differences, and confirmed that ^18^F-FCH uptake correlates with other markers of BAT such as adipose tissue radiodensity in a larger clinical cohort. This widely available clinical PET radiotracer may offer an alternative method to quantify human BAT in patients at room temperature. We have also determined the phosphatidylcholine species synthesized from choline in human brown adipocytes, but further work is required to determine how these phospholipids regulate BAT function.

## 4. MATERIALS AND METHODS

### 4.1 ^18^F-FCH uptake by BAT *in vivo* in healthy volunteers

Six healthy male volunteers were recruited to a randomised crossover study examining the effect of warm and cold exposure on ^18^F-fluorocholine (^18^F-FCH) uptake by BAT. As no previous studies have investigated ^18^F-FCH to detect BAT, our previous ^18^F-FDG data in BAT and WAT in similar individuals was used to determine that 6 participants provided >80% power to detect a difference in ^18^F-FCH standardised uptake values (SUV) of 0.5 g/mL between BAT and WAT depots [23, 25, 48]. Inclusion criteria were as follows: male; body mass index (BMI) 18.5–25[Kg/[m^2^; no acute or chronic medical conditions; on no regular medications; no claustrophobia; no contraindication to MRI scan (e.g. cochlear implant, pacemaker); alcohol intake ≤ 14 units/ week; screening blood tests within an acceptable range (blood count, renal function, liver function, thyroid function and random glucose); ability to provide informed consent; weight change of less than 5% in the preceding 6 months. While participants could not be blinded to the ambient room temperature, the analysis of the PET/MR scans and blood samples were performed by researchers blinded to the order of the warm and cold visits. Approval was obtained from the South East Scotland Research Ethics Committee (ethics number 17/SS/0095) as was informed consent from each volunteer, and all research was performed in accordance with the Declaration of Helsinki. The protocol and data from visit 1 have been presented previously [12].

Following recruitment, volunteers attended the Edinburgh Clinical Research Facility (ECRF) at the Royal Infirmary of Edinburgh and Edinburgh Imaging Facility (EIF) in the Queen’s Medical Research Institute on 3 further occasions (Figure 1A). Volunteers attended after overnight fast after avoiding alcohol and exercise for 2 days prior to each visit. Each study visit was separated by at least 2 weeks. At study visit 1, fasting blood samples were obtained for measurement of glucose, insulin and NEFAs. At T[=[0[minutes, participants were placed in a room cooled to 16–17 °C (cold room) for 2 h to activate BAT. Participants were checked every 15[minutes for signs or symptoms of shivering. After 1 hour of cold exposure, volunteers received an intravenous injection of 75[MBq of ^18^F-FDG and underwent a PET/MRI scan to quantify BAT mass and activity 1 hour later. Following the scan, volunteers returned home. All participants had detectable ^18^F-FDG uptake by BAT so proceeded to the final two visits of the study protocol.

Prior to study visit 2, participants were randomised to undergo ^18^F-FCH PET/MR scanning during either warm (∼24°C) or cold conditions (16-17°C) in a crossover design. Apart from room temperature, study visits 2 and 3 were identical. Volunteers were placed in the warm or cold room for 2 hours, fasting blood samples were obtained every 15 minutes during the first hour. Blood samples were not obtained in one volunteer due to technical difficulties, so data are presented for the remaining 5 participants. Participants were checked every 15[minutes for signs or symptoms of shivering. After 1 hour, participants received an intravenous injection of the radiotracer ^18^F-FCH (150MBq) and a PET/MR scan was performed 1 hour later.

#### 4.1.1 PET/MR scanning protocol

Full details have been published previously [23]. All participants received an IV bolus of ^18^F-FDG (75MBq) 1 hour before being placed supine on a Siemens mMR scanner (Siemens Healthineers). Following initial localization, a standard MRAC_GRAPPA scan was acquired for each bed position, used to calculate a standard umap for attenuation correction. A three-dimensional T1-weighted Dixon VIBE acquisition was used to generate images at 1.34 and 2.56[ms (repetition time (TR)[=[4.02) to calculate a fat fraction map (FFM).

#### 4.1.2 PET/MR image analysis

^18^F-FDG and ^18^F-FCH uptake by BAT were quantified using Analyze 12.0 (AnalyzeDirect, Overland Park, KS) as previously described [23]. Following registration, the MRI fat and water images were used to generate a FFM of each participant using the following equation:

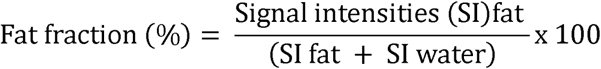

A median spatial filter was applied to the FFM, voxels with a fat fraction <50% were removed to generate a fat fraction region of interest (ROI). For the ^18^F-FDG analysis, the registered PET images for each participant were thresholded using a standard uptake value (SUV)_lean_ of ≥1.2 g/[mL [13]. Any residual ROIs that encompassed the brain were removed manually, and ROIs for analysis were composed of voxels meeting both the FFM and PET thresholds. Finally, to remove PET blooming and boundary artefacts image erosion of the BAT ROIs was performed using a 3[×[3[×[1 structuring element. Data are presented for the BAT SUV_mean_ and BAT volume. ROIs were also drawn manually around subcutaneous abdominal WAT, ensuring they were identically sized between volunteers and between each visit.

For the ^18^F-FCH analysis, initially the BAT and WAT ROIs drawn using the ^18^F-FDG-PET data were co-registered to the ^18^F-FCH-PET scans to allow quantification of ^18^F-FCH SUV_mean_ for both BAT and WAT. Thereafter, ^18^F-FCH uptake by BAT was quantified without the use of the ^18^F-FDG ROIs. The above process was repeated, except the ^18^F-FCH PET images were thresholded using a SUV of ≥0.5 g/[mL.

#### 4.1.3 Biochemical assays

Plasma glucose (Sigma-Aldrich) and NEFAs (Wako Diagnostics) were measured using colorimetric kits, with serum insulin (Mercodia) and plasma noradrenaline (LDN) by enzyme-linked immunosorbent assay.

### 4.2 ^18^F-FCH uptake by supraclavicular adipose tissue in patients undergoing PET/CT

A retrospective analysis was performed of patients who had undergone ^18^F-FCH PET/CT scanning at the EIF between June 2015 and May 2021 as part of their clinical care (Table S1). Local Caldicott guardian approval was obtained for this study, and scans were anonymised by the clinical team prior to analysis by the research team. Patients received either 4 MBq/ Kg (prostate) or 2 MBq/ Kg (parathyroid) of ^18^F-FCH (up to a maximum of 300 MBq) intravenously 1 hour before being placed supine in a hybrid PET/CT scanner. Two models of hybrid PET–CT scanners were used, a Biograph mCT (Siemens Medical Systems) or a Discovery 710 (GE Healthcare). Scans were performed at room temperature (∼20-21°C). All patients underwent a CT scan for attenuation correction (non-enhanced, 120kV) with tube modulation applied (50 mAs) followed by static PET of the upper body using 4-minute beds. On the Biograph, images were reconstructed using Siemens UltraHD reconstruction (2 iterations and 21 subsets). On the Discovery, the QClear reconstruction was used with a beta value of 400. Images were analysed using Analyze 12.0 (AnalyzeDirect, Overland Park, KS).

CT images were thresholded between the radiodensity range of −190 to −10 Hounsfield Units, and ROIs were drawn around supraclavicular adipose tissue, a typical BAT depot [49]. The data presented are the mean SUV in these ROIs.

### 4.3 Human adipose tissue collections

Male and female euthyroid participants were recruited (Table S4) who were attending NHS Lothian for either: 1) elective thyroid or parathyroid surgery or 2) elective nephrectomy from living kidney donors. Local ethical approval was obtained (research ethics committee numbers 15/ES/0094, 20/ES/0061) as was consent from each participant. For neck patients, adipose tissue was obtained intra-operatively from the neck as previously described [48], both deep to the lateral thyroid lobe either adjacent to the longus colli muscle or to the oesophagus (BAT), and more superficially from the subcutaneous tissue (WAT). Tissue collections were performed by one surgeon. For living donor nephrectomy patients, adipose tissue was obtained from the upper (BAT) and lower (WAT) poles of the kidney, and from the subcutaneous fat at the incision site (WAT). Tissue samples were either immediately frozen at −80°C for qPCR (both neck and peri-renal samples), or the stromal vascular fraction was isolated and cultured for qPCR and the ^15^N-choline tracing experiments detailed below (neck AT only).

Cells were cultured as described previously [48]. In brief, adipose tissue was incubated in Krebs–Henseleit buffer containing 0.2% collagenase type 1 for 45 minutes at 37°C. Following centrifugation at 800g at room temperature for 10 minutes, the pellet was resuspended and passed through a 100μm filter. Samples were subject to centrifugation at 200g for 5 minutes and the pellet resuspended in DMEM containing 10% fetal bovine serum (FBS) and plated in 6-well plates for culture. Following proliferation and passage, cells were differentiated in DMEM containing 10% FBS with the addition of 20nM insulin, 1nM tri-iodothyronine, 500μM IBMX, 125μM indomethacin and 500nM dexamethasone for 7 days. Differentiated pre-adipocytes were then cultured in DMEM medium containing 10% FBS, 1nM tri-iodothyronine and 20nM insulin for a further 7 days prior to experiments.

#### 4.3.1 Quantitative real time PCR

qPCR was performed as previously described [23]. Adipose tissue was homogenized in QIAzol using a TissueLyser (Qiagen). mRNA was extracted from tissue and cells using the RNeasy Lipid Kit (Qiagen) and complementary DNA generated using the Qiagen QuantiTect reverse transcription kit. qPCR was performed in triplicate using a Roche Lightcycler 480, using gene-specific primers (Invitrogen) and fluorescent probes from the Roche Universal Probe Library, or with Taqman assays (Table S5). Transcript levels are presented as the ratio of the abundance of the gene of interest: mean of abundance of the control genes *PPIA* and *RNA18S5*.

### 4.4 ^15^N-choline utilisation by primary brown adipocytes

Human brown adipocytes were incubated with either vehicle or 400µM [^15^N]-choline chloride (Sigma), in the presence of either vehicle or 10µM L-noradrenaline bitartrate salt monohydrate (Sigma) for 24 hours in a 2 x 2 design in triplicate. Following incubation, the medium was removed, wells were washed with ice cold DPBS, cells were scraped in ice-cold isopropanol and the extracts stored at –80°C until analysis.

#### 4.4.1 Nano Lipidomics

Lipid metabolites from brown adipocytes were analysed using a nano-LC-MS/MS methodology [50]. Lipids were extracted from cell pellets in 100% isopropanol (MS grade) and extracts further cleared by centrifugation. 1 μL of lipid extract was loaded onto an Aurora C18 column (IonOptics, Australia), attached to a Thermo Ultimate 3000 RSLCnano HPLC. Flow rate was 300 nL/min and column temperature was 50°C. The column was equilibrated in 70% buffer A (60% acetonitrile, 0.1% formic acid, 10mM ammonium formate) and 30% buffer B (90% isopropanol, 10% acetonitrile, 0.1% formic acid, 10mM ammonium formate) and the following gradient was applied (time/%B): 1/1, 4/30, 5/35, 8/51, 20/61, 25/70, 30/99. Lipids were eluted into a Fusion Lumos mass spectrometer (Thermo Scientific) in positive mode with a scan range of 200-1600 in MS1 and resolution 500,000. Data-dependent MS2 scans were acquired in the ion trap (stepped collision energies 20, 30, 40; Rapid scan rate, scan range 120-1200). Other settings were as standard.

The nano-LC-MS data were first centroided and converted to MZML using MSConvert (http://proteowizard.sourceforge.net/). The centroided data were deconvolved with XCMS online (https://xcmsonline.scripps.edu/) [51]: Feature detection; method – CentWave; mass error 3ppm, minimum and maximum peak width 3 and 30 s respectively, mzdiff 0.01, S/N threshold 6, integration method 1, prefilter peaks 3, prefilter intensity 100, noise filter 0: RT (retention time) correction; method – Obiwarp, profstep 1: Alignment; minfrac 0.5, *m/z* width 0.015, bw 5, min samp 1, max samp 100: Annotation; Search for isotopes+adducts, mz absolute error 0.015, ppm error 5. XCMS generates a Microsoft Excel based XY matrix containing paired RT and *m/z*, along with the extracted ion chromatogram (EIC) area for each profiled sample.

Applying PutMedID operated within Taverna Workbench 1.7.2 [52, 53], peak to peak Pearson correlations (workflow 1) were calculated (+/− 5 s), peaks that show high levels of correlation (0.8>) are grouped as *m/z* features associated with the same compound (i.e. an *m/z* group). Workflow 2 next calculates accurate mass differences between ions in each *m/z* group and applies known values of accurate mass difference for commonly formed adducts, isotopes and in-source fragments, to enable the annotation of the parent *m/z* as well as the various adduct and isotope ions. Finally, the neutral accurate mass is calculated and in turn matched to possible molecular formula(s) (<5ppm). Workflow 3 matches compound molecular formula to metabolite name(s) based upon the Manchester Metabolomics Database (MMD: http://dbkgroup.org/MMD/).

Particular attention was placed upon the annotation of lipid species. Non-labelled species were quickly ascertained through the typical isotope spacing (+1.00335) for single charged ions, whereas N15 labelled species were quickly ascertained through isotopic spacing of the N14 to N15 incorporated lipid (+0.997035). Matching monosiotopic (m+0) and (m+1) ions were identified, with the latter comprising combined intensities of 13C1– and 15N1- containing ions. The theoretical natural abundance of 13C1 ions, based upon number of carbons was subtracted and the remaining (m+1) peak areas were used to obtain a (m+1)/(m+0) labelling ratio. 15N-labelled phosphatidylcholines were identified by clear labelling in the choline samples and near (or complete) absence of labelling in the unlabelled samples. Data was minimised to only include H+ adducts (Table S3).

### 4.5 Statistical Analyses

All statistical analyses were performed using Prism software (GraphPad version 10.2.3, USA except lipidomic data which were analysed as described above). All data were analysed for normal distribution within each experimental group using the Shapiro-Wilk test. Normally distributed data were analysed by ANOVA or *t* tests, as appropriate. Where data were not normally distributed, non-parametric tests were used, the specific statistical tests used are detailed in the respective figure captions. All data points are presented, with the mean[±[SEM / SD as appropriate. A P value[<[0.05 was considered statistically significant. Figure legends describe the number of biologically independent samples and the specific statistical tests used. All data generated or analysed during this study are included in this published article.

## Abbreviations

^18^F-FCH: 18F-Fluorocholine
^18^F-FDG: ^18^F-fluorodeoxyglucose
AT: adipose tissue
BMI: body mass index
BAT: brown adipose tissue
CVD: cardiovascular disease
LPCs: lysophosphatidylcholines
NADR: noradrenaline
PCs: phosphatidylcholines
PET: positron emission tomography
ROIs: regions of interest
T2DM: type 2 diabetes mellitus
UCP1: uncoupling protein 1
WAT: white adipose tissue

## ACKNOWLEDGEMENTS

This work was supported by grants from the Medical Research Council (MR/S035761/1, MR/W01937X/1), the Chief Scientist Office (SCAF/17/02), and the Wellcome Trust Institutional Strategic Support Fund (IS3-R42) to R.H.S, and a Wellcome Trust multiuser equipment grant (208402/Z/17/Z) to A.V.K. and A.J.F. We thank the nurses and other staff at the Edinburgh Clinical Research Facility, and Tim Clark and the radiographers at the Edinburgh Imaging Facility QMRI for their assistance. We thank Ruth Andrew and the Scottish Metabolomics Network for academic facilitation.

## Author contributions

Conceptualization, R.H.S; methodology, K.J.S., L.E.R., G.R.B, C.G., A.V.K., M.G., A.M.F., S.J.W., J.D.T., E.V.B., D.P., A.J.F., R.H.S.; Investigation – K.J.S., L.E.R., L.D.B., T.C.K., G.R.B, A.K., C.G., G.C.O., N.Z.M.H., J.D.T., W.A., S.J.W., A.J.F., R.H.S.; Formal analyses, K.J.S., L.E.R., T.C.K., W.A., A.J.F., R.H.S.; Writing – original draft, K.J.S and R.H.S; Writing – review & editing, K.J.S., L.E.R., L.D.B., T.C.K., G.R.B, A.K., C.G., A.V.K., M.G., G.C.O., A.M.F., N.Z.M.H., J.D.T., W.A., S.J.W., E.V.B., D.P., A.J.F., R.H.S.. Visualization, K.J.S., R.H.S.; R.H.S. supervised the study and acquired the funding.

## Declaration of interests

The authors declare no competing interests.

## Data availability

Data will be made available upon request.

## Notes

### Competing Interest Statement

The authors have declared no competing interest.

