## Supplementary material for "Human brown adipose tissue demonstrates substantial ^18^F-fluorocholine uptake at room temperature for phosphatidylcholine synthesis": Figure S1

**SUPPLEMENTAL INFORMATION**

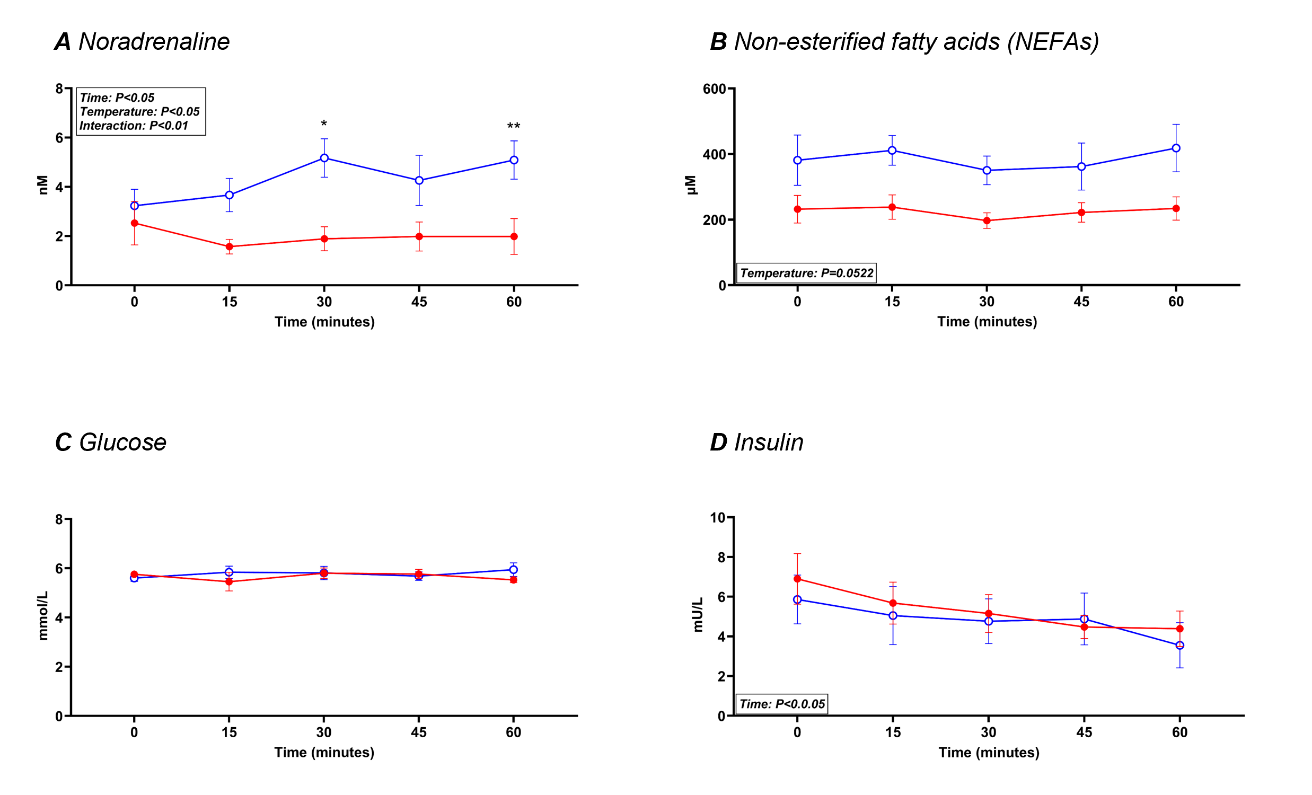
***Figure S1	Effect of cold exposure on biochemical measurements in healthy volunteers***

*Data are mean ± SEM for circulating A) noradrenaline, B) non-esterified fatty acids (NEFAs), C) glucose and D) insulin in the healthy volunteers (all n=5) exposed to either warm (~24°C, red closed circles) or cold (~16°C, blue open circles) conditions. Cold exposure increased plasma noradrenaline and serum NEFAs. Data were analysed by 2-way repeated measures ANOVA with Sidak’s multiple comparison post-hoc testing.*

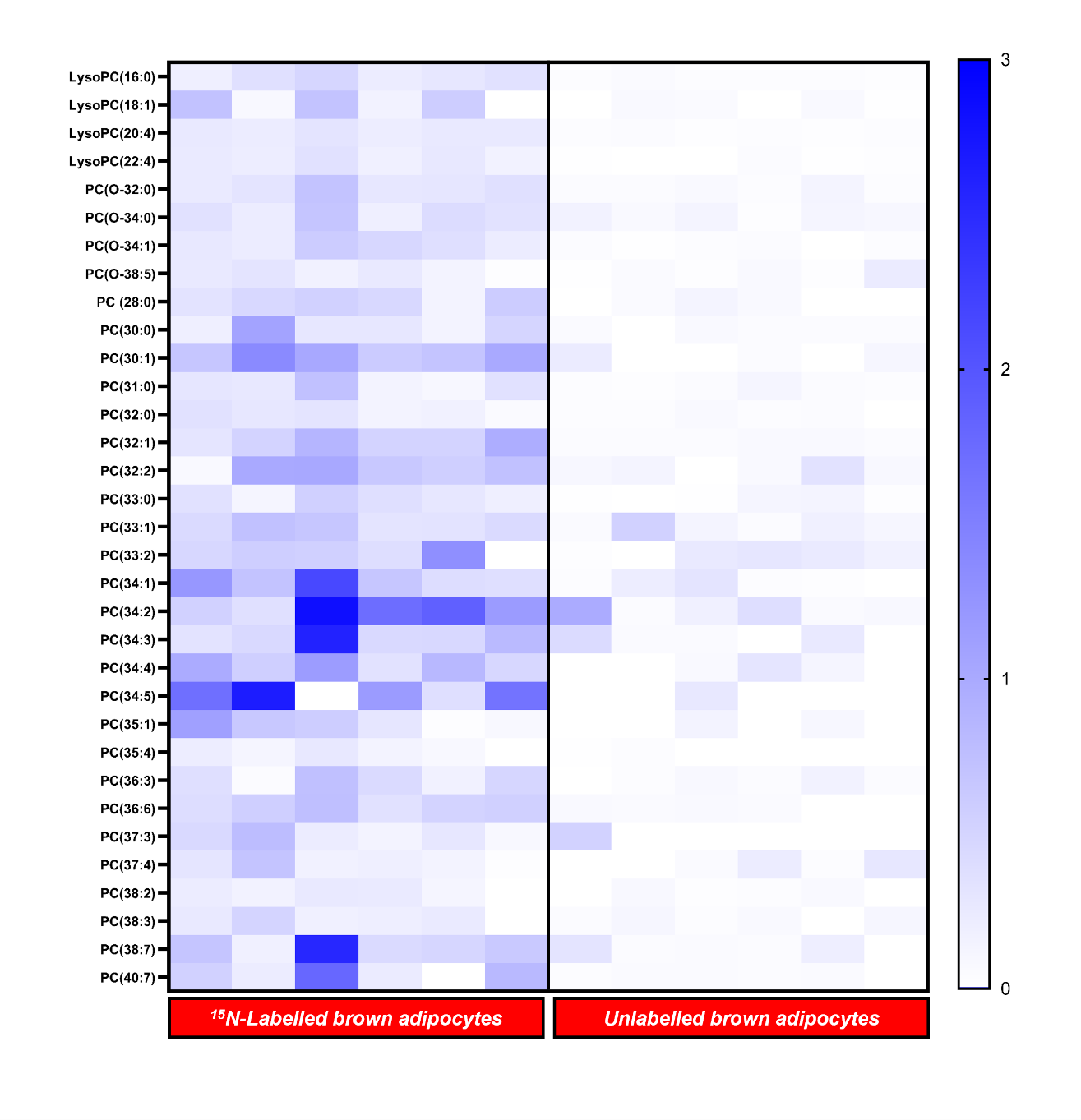
***Figure S2	PCs and LPCs identified from ^15^N-Choline-labelled and unlabelled human brown adipocytes***

*Lipidomics was performed on human brown adipocytes incubated with either 400µM ^15^N-choline or vehicle for 24 hours. Heat map detailing validation of m+1 (^15^N) containing phosphatidylcholine (PC) and lyso-PC (LPC) species, showing labelled vs unlabelled cells. Values presented are the ratios of (m+1)/(m+0), following theoretical subtraction of the contribution of (m+1) from the natural abundance of ^13^C.*

***Table S1	Patient data from ^18^F-FCH PET/CT analysis of clinical scans***

*Data are mean ± SD. Scans were performed between June 2015 and May 2021.*

| Number of patients | 76 |
| --- | --- |
| Gender | 26 Female, 50 Male |
| Number with prostate cancer | 41 |
| Number with primary hyperparathyroidism | 35 |
| Age (y) | 58.9 ± 14.0 |
| Weight (Kg) | 83.6 ± 15.4 |
| Height (m) | 1.73 ± 0.10 |
| BMI (Kg/m^2^) | 27.9 ± 4.6 |

***Table S2	Expression of genes involved in choline metabolism in human adipocytes***

| **Gene** | **Mean Brown** | **Mean White** | **P value (adjusted)** |
| --- | --- | --- | --- |
| **Kennedy** | | | |
| *CHKA* | 29.78 | 27.33 | 0.99997 |
| *CHKB* | 66.44 | 62.74 | 0.99997 |
| *PCYT1A* | 118.45 | 122.32 | 0.99997 |
| *CHPT1* | 113.08 | 120.63 | 0.99997 |
| *CEPT1* | 224.44 | 191.98 | 0.98436 |
| **Lands** | | | |
| *LCAT* | 19.11 | 19.45 | 0.99997 |
| *LPCAT1* | 52.24 | 46.84 | 0.99997 |
| *LPCAT2* | 24.65 | 25.23 | 0.99997 |
| *LPCAT3* | 109.70 | 69.81 | 0.82822 |
| *LPCAT4* | 30.91 | 22.60 | 0.99997 |
| *PLA2G2A* | 69.19 | 73.71 | 0.99997 |
| *PLA2G4A* | 90.14 | 38.36 | **0.00098** |
| *PLA2G4B* | 4.53 | 4.18 | 0.99997 |
| *PLA2G4C* | 33.99 | 29.17 | 0.99997 |
| *PLA2G5* | 12.80 | 13.17 | 0.99997 |
| *PLA2G6* | 23.82 | 23.47 | 0.99997 |
| *PLA2G15* | 31.12 | 27.44 | 0.99997 |
| *PLA2G16* | 351.97 | 234.28 | 0.43144 |
| **Betaine** | | | |
| *AHCY* | 305.06 | 320.48 | 0.99997 |
| *ALDH7A1* | 96.52 | 130.86 | 0.74466 |
| *BHMT2* | 76.32 | 67.93 | 0.99997 |
| *CHDH* | 2.26 | 1.96 | 0.99997 |
| *MAT2A* | 144.84 | 120.58 | 0.99997 |
| *MAT2B* | 238.40 | 237.07 | 0.99997 |
| *PEMT* | 360.23 | 386.60 | 0.99997 |

***Table S3	PC and LPC species incorporating ^15^N-choline quantified in human brown adipocytes***

*Data showing incorporation of ^15^N-choline into selected phosphatidylcholines (PCs) and lysophosphatidylcholines (LPCs), as measured by LC-MS. Cells were either unlabelled or incubated with ^15^N-choline in the presence of either vehicle or 10µM noradrenaline for 24h. Values presented are the ratios of (m+1)/(m+0), following theoretical subtraction of the contribution of (m+1) from the natural abundance of ^13^C (see methods). Negative values are depicted as zero.*

|  | **^15^N-Choline Vehicle** | | | **^15^N-Choline Noradrenaline** | | | **Unlabelled Vehicle** | | | **Unlabelled Noradrenaline** | | |
| --- | --- | --- | --- | --- | --- | --- | --- | --- | --- | --- | --- | --- |
|  | **P1** | **P2** | **P3** | **P1** | **P2** | **P3** | **P1** | **P2** | **P3** | **P1** | **P2** | **P3** |
| **LysoPC(16:0)** | 0.185 | 0.354 | 0.479 | 0.220 | 0.275 | 0.351 | 0.035 | 0.058 | 0.026 | 0.029 | 0.029 | 0.014 |
| **LysoPC(18:1)** | 0.709 | 0.072 | 0.701 | 0.147 | 0.588 | 0.000 | 0 | 0.065 | 0.051 | 0 | 0.059 | 0.004 |
| **LysoPC(20:4)** | 0.254 | 0.219 | 0.310 | 0.209 | 0.253 | 0.247 | 0.031 | 0.035 | 0.023 | 0.032 | 0.023 | 0.030 |
| **LysoPC(22:4)** | 0.243 | 0.212 | 0.346 | 0.166 | 0.267 | 0.153 | 0.012 | 0 | 0 | 0.037 | 0.000 | 0.015 |
| **PC(O-32:0)** | 0.243 | 0.314 | 0.705 | 0.274 | 0.289 | 0.362 | 0.040 | 0.043 | 0.062 | 0.032 | 0.129 | 0.033 |
| **PC(O-34:0)** | 0.346 | 0.221 | 0.679 | 0.185 | 0.403 | 0.338 | 0.144 | 0.064 | 0.123 | 0.022 | 0.115 | 0.087 |
| **PC(O-34:1)** | 0.263 | 0.218 | 0.598 | 0.466 | 0.375 | 0.213 | 0.039 | 0.001 | 0.026 | 0.040 | 0 | 0.026 |
| **PC(O-38:5)** | 0.254 | 0.312 | 0.159 | 0.252 | 0.132 | 0.019 | 0.008 | 0.050 | 0.022 | 0.070 | 0.023 | 0.225 |
| **PC (28:0)** | 0.319 | 0.453 | 0.530 | 0.452 | 0.135 | 0.589 | 0 | 0.050 | 0.129 | 0.069 | 0 | 0 |
| **PC(30:0)** | 0.181 | 1.075 | 0.273 | 0.276 | 0.133 | 0.487 | 0.056 | 0 | 0.062 | 0.045 | 0.042 | 0.046 |
| **PC(30:1)** | 0.662 | 1.365 | 1.020 | 0.607 | 0.691 | 1.003 | 0.233 | 0 | 0 | 0.041 | 0 | 0.096 |
| **PC(31:0)** | 0.284 | 0.259 | 0.718 | 0.134 | 0.085 | 0.343 | 0.035 | 0.013 | 0.036 | 0.117 | 0.042 | 0.028 |
| **PC(32:0)** | 0.352 | 0.273 | 0.312 | 0.137 | 0.159 | 0.054 | 0.033 | 0.033 | 0.068 | 0.030 | 0.038 | 0 |
| **PC(32:1)** | 0.305 | 0.514 | 0.861 | 0.508 | 0.516 | 0.959 | 0.041 | 0.040 | 0.042 | 0.063 | 0.059 | 0.038 |
| **PC(32:2)** | 0.067 | 1.012 | 1.023 | 0.647 | 0.561 | 0.729 | 0.092 | 0.127 | 0 | 0.069 | 0.335 | 0.073 |
| **PC(33:0)** | 0.347 | 0.109 | 0.545 | 0.371 | 0.271 | 0.184 | 0.008 | 0 | 0.010 | 0.114 | 0.118 | 0.021 |
| **PC(33:1)** | 0.426 | 0.719 | 0.666 | 0.314 | 0.329 | 0.428 | 0.056 | 0.535 | 0.128 | 0.044 | 0.167 | 0.105 |
| **PC(33:2)** | 0.466 | 0.575 | 0.543 | 0.381 | 1.313 | 0.000 | 0.015 | 0 | 0.258 | 0.288 | 0.242 | 0.159 |
| **PC(34:1)** | 1.217 | 0.700 | 2.147 | 0.667 | 0.398 | 0.376 | 0.030 | 0.211 | 0.306 | 0.034 | 0.023 | 0 |
| **PC(34:2)** | 0.532 | 0.354 | 2.826 | 1.708 | 1.867 | 1.174 | 0.981 | 0.040 | 0.174 | 0.377 | 0.052 | 0.074 |
| **PC(34:3)** | 0.326 | 0.445 | 2.599 | 0.446 | 0.453 | 0.808 | 0.421 | 0.049 | 0.058 | 0 | 0.258 | 0 |
| **PC(34:4)** | 0.978 | 0.564 | 1.164 | 0.340 | 0.831 | 0.465 | 0 | 0 | 0.064 | 0.296 | 0.115 | 0 |
| **PC(34:5)** | 1.691 | 2.663 | 4.569 | 1.166 | 0.371 | 1.647 | 0 | 0 | 0.264 | 0 | 0 | 0 |
| **PC(35:1)** | 1.107 | 0.640 | 0.586 | 0.285 | 0.015 | 0.077 | 0 | 0 | 0.138 | 0 | 0.091 | 0 |
| **PC(35:4)** | 0.206 | 0.114 | 0.269 | 0.138 | 0.082 | 0 | 0.009 | 0.026 | 0 | 0 | 0 | 0 |
| **PC(36:3)** | 0.374 | 0.036 | 0.737 | 0.426 | 0.161 | 0.476 | 0 | 0.034 | 0.081 | 0.039 | 0.150 | 0.047 |
| **PC(36:6)** | 0.395 | 0.560 | 0.746 | 0.344 | 0.507 | 0.547 | 0.067 | 0.047 | 0.060 | 0.050 | 0 | 0 |
| **PC(37:3)** | 0.447 | 0.767 | 0.214 | 0.135 | 0.271 | 0.081 | 0.528 | 0 | 0 | 0 | 0 | 0 |
| **PC(37:4)** | 0.296 | 0.689 | 0.154 | 0.187 | 0.140 | 0.021 | 0 | 0 | 0.050 | 0.213 | 0.032 | 0.276 |
| **PC(38:2)** | 0.215 | 0.149 | 0.255 | 0.243 | 0.117 | 0.000 | 0 | 0.075 | 0.023 | 0.032 | 0.060 | 0 |
| **PC(38:3)** | 0.251 | 0.503 | 0.175 | 0.199 | 0.238 | 0 | 0.044 | 0.104 | 0.035 | 0.059 | 0 | 0.103 |
| **PC(38:7)** | 0.678 | 0.168 | 2.533 | 0.427 | 0.472 | 0.625 | 0.307 | 0.045 | 0.049 | 0.040 | 0.189 | 0 |
| **PC(40:7)** | 0.538 | 0.217 | 1.782 | 0.228 | 0 | 0.822 | 0.032 | 0.050 | 0.047 | 0.046 | 0.053 | 0 |

***Table S4	Patient details for adipose tissue collections***

| Experiment/ Subject number | Operation | Underlying diagnosis | Age (years) | Sex | BMI (Kg/m^2^) |
| --- | --- | --- | --- | --- | --- |
| Adipocytes for qPCR |  |  |  |  |  |
| 1 | Parathyroidectomy | Primary hyperparathyroidism | 51 | Female | 26.3 |
| 2 | Thyroid lobectomy | Multinodular goitre | 41 | Female | 28.6 |
| 3 | Thyroidectomy | Papillary microcarcinoma | 35 | Female | 37.0 |
| 4 | Parathyroidectomy | Primary hyperparathyroidism | 70 | Female | 32.5 |
| 5 | Parathyroidectomy | Primary hyperparathyroidism | 58 | Female | 22.9 |
| 6 | Parathyroidectomy | Primary hyperparathyroidism | 59 | Female | 24.4 |
| 7 | Thyroid lobectomy | Papillary thyroid carcinoma | 36 | Female | 28.0 |
| 8 | Parathyroidectomy | Primary hyperparathyroidism | 61 | Female | 23.3 |
| 9 | Parathyroidectomy | Primary hyperparathyroidism | 67 | Female | 18.7 |
| 10 | Parathyroidectomy | Primary hyperparathyroidism | 62 | Female | 30.8 |
| 11 | Parathyroidectomy | Primary hyperparathyroidism | 63 | Female | 33.9 |
| 12 | Parathyroidectomy | Primary hyperparathyroidism | 68 | Female | 21.9 |
| Supraclavicular adipose tissue |  |  |  |  |  |
| 1 | Parathyroidectomy | Primary hyperparathyroidism | 67 | Female | 27.9 |
| 2 | Parathyroidectomy | Primary hyperparathyroidism | 65 | Female | 29.7 |
| 3 | Parathyroidectomy | Primary hyperparathyroidism | 37 | Male | 26.4 |
| 4 | Parathyroidectomy | Primary hyperparathyroidism | 59 | Female | 26.7 |
| 5 | Parathyroidectomy | Primary hyperparathyroidism | 54 | Male | 31.1 |
| 6 | Parathyroidectomy | Primary hyperparathyroidism | 54 | Female | 34.1 |
| 7 | Thyroidectomy | Papillary thyroid carcinoma | 36 | Female | 37.9 |
| 8 | Parathyroidectomy | Primary hyperparathyroidism | 65 | Male | 31.1 |
| Peri-renal adipose tissue |  |  |  |  |  |
| 1 | Nephrectomy | Healthy kidney donor | 67 | Female | 22.9 |
| 2 | Nephrectomy | Healthy kidney donor | 53 | Male | 24.4 |
| 3 | Nephrectomy | Healthy kidney donor | 48 | Male | 28.0 |
| 4 | Nephrectomy | Healthy kidney donor | 53 | Female | 23.3 |
| 5 | Nephrectomy | Healthy kidney donor | 47 | Female | 18.7 |
| 6 | Nephrectomy | Healthy kidney donor | 54 | Female | 30.8 |
| 7 | Nephrectomy | Healthy kidney donor | 37 | Male | 33.9 |
| ^15^N-Choline labelling |  |  |  |  |  |
| 1 | Thyroidectomy | Papillary microcarcinoma | 35 | Female | 37.0 |
| 2 | Parathyroidectomy | Primary hyperparathyroidism | 60 | Female | 30.6 |
| 3 | Parathyroidectomy | Primary hyperparathyroidism | 61 | Female | 23.3 |

***Table S5	Primer sequences for qPCR and corresponding probe numbers***

*Assays were either performed using the following primer sequences and the relevant Roche probe library or using the following Taqman assay ID numbers.*

| Gene Name | Primer sequences 5’ to 3’ | Roche UPL Probe number/ Taqman ID number |
| --- | --- | --- |
| *PPIA*  (cyclophilin A) | F: atgctggacccaacacaat | 48 |
|  | R: tctttcactttgccaaacacc |  |
| *RNA18S5*  (18S) | F: cttccacaggaggcctacac | 46 |
|  | R: cgcaaaatatgctggaacttt |  |
| *UCP1* | F: ctcaccgcagggaaagaa | 25 |
|  | R: ggttgcccaatgaatactgc |  |
| *SLC5A7* | F: N/A | Hs00222367_m1 |
|  | R: N/A |  |
| *SLC22A1* | F: N/A | Hs00427552_m1 |
|  | R: N/A |  |
| SLC22A2 | F: N/A | Hs01010726_m1 |
|  | R: N/A |  |
| *SLC25A48* | F: N/A | Hs00415075_m1 |
|  | R: N/A |  |
| *SLC44A1* | F: N/A | Hs00223114_m1 |
|  | R: N/A |  |
| *SLC44A2* | F: N/A | Hs01105936_m1 |
|  | R: N/A |  |
| *SLC44A3* | F: N/A | Hs00537043_m1 |
|  | R: N/A |  |
